# Regulation of nuclear actin levels and MRTF/SRF target gene expression during PC6.3 cell differentiation

**DOI:** 10.1101/2022.06.14.496089

**Authors:** Salla Kyheröinen, Alise Hyrskyluoto, Maria Sokolova, Maria K. Vartiainen

## Abstract

Actin has important functions in both cytoplasm and nucleus of the cell, with active nuclear transport mechanisms maintaining the cellular actin balance. Nuclear actin levels are subject to regulation during many cellular processes from cell differentiation to cancer. Here we show that nuclear actin levels increase upon differentiation of PC6.3 cells towards neuron-like cells. Photobleaching experiments demonstrate that this increase is due to decreased nuclear export of actin during cell differentiation. Increased nuclear actin levels lead to decreased nuclear localization of MRTF-A, a well-established transcription cofactor of SRF. In line with MRTF-A localization, transcriptomics analysis reveals that MRTF/SRF target gene expression is first transiently activated, but then substantially downregulated during PC6.3 cell differentiation. This study therefore describes a novel cellular context, where regulation of nuclear actin is utilized to tune MRTF/SRF target gene expression during cell differentiation.

## Introduction

Actin is a protein widely known from its essential cytoplasmic functions, providing both structure, and together with motor protein myosin, motility to cells. These functions are largely based on the ability of actin to polymerize from monomers (G-actin) to helical filaments (F-actin) (Pollard & Cooper, 2009). More recently, actin has also emerged as an important nuclear protein, with functional roles in many essential nuclear processes from gene expression to maintenance of genomic integrity (Hyrskyluoto & Vartiainen, 2020). Some of these nuclear functions of actin are dependent on polymerization. For example, directing DNA double strand breaks (DSBs) towards safe nuclear compartments for repair requires nuclear actin polymerization regulated by different actin-binding proteins (ABPs) (Caridi et al., 2018; Schrank et al., 2018). Similarly, nuclear expansion and chromatin decondensation after cell division (Baarlink et al., 2017), as well as loading of replication factors and initiation of DNA replication (Parisis et al., 2017), depend on actin dynamics. Interestingly, also actin monomers have functional roles within the nucleus as part of several chromatin remodeling and modifying complexes (Klages-Mundt et al., 2018). Actin also associates with the transcription machinery all the way from regulating the activity of specific transcription factors (Vartiainen et al., 2007 and see below), transcription initiation (Hofmann et al., 2004; Sokolova et al., 2018) to RNA polymerase II pause-release (Qi et al., 2011), pre-mRNA processing (Viita et al., 2019) and RNA polymerase clustering (Wei et al., 2020).

Considering the essential functions of actin in both nucleus and cytoplasm, cells must ensure appropriate regulation of protein flux between these compartments. Indeed, the nuclear and cytoplasmic pools of actin are dynamically connected, and actin shuttles rapidly in and out of the nucleus by active transport. Nuclear import of actin requires Importin-9 (Ipo9) together with the small ABP cofilin (Dopie et al., 2012), while export is carried out by Exportin-6 (Exp6) and ABP profilin (Dopie et al., 2012; Stuven et al., 2003). One of the limiting factors for nuclear import and export rates is the availability of actin monomers (Dopie et al., 2012; Skarp et al., 2013), which depends on actin polymerization, the number of binding events with ABPs and association to larger molecular machineries, such as the chromatin remodelers. Thus, the nucleo-cytoplasmic shuttling process of actin offers several regulatory points to tune the cellular actin balance, and alterations in nuclear actin levels have been reported in several conditions. In mammary epithelial cells, extracellular matrix component laminin-111 or lack of growth factors decrease nuclear actin levels and reduce transcription (Spencer et al., 2011). Here laminin-111 attenuates the PI3-kinase pathway, which results in Exp6 upregulation and thus regulation of nuclear actin at the level of enhanced nuclear export. Defects in this mechanism are observed in malignant cells (Fiore et al., 2017). Nuclear actin levels are also reduced in epidermal stem cells in response to mechanical stress, with functional implications in heterochromatin anchoring, transcription and Polycomb-mediated gene silencing. Accumulation of emerin at the outer nuclear membrane and enrichment of non-muscle myosin IIA cause local actin polymerization, which decreases the available actin monomers. Hence, in this cell model, nuclear actin levels are regulated at the level of nuclear import (Le et al., 2016). On the other hand, *Xenopus* oocytes contain massive amounts of nuclear actin, which forms a filamentous mesh required to support the structure of these huge nuclei (Bohnsack et al., 2006), and protect for example ribonucleoprotein droplets against gravity (Feric & Brangwynne, 2013). In these cells, nuclear actin levels seem to be regulated at the level of export, via post-transcriptional silencing of Exp6 (Bohnsack et al., 2006) via an unknown mechanism. Increased nuclear actin levels have also been reported during differentiation of HL-60 cells towards macrophages upon PMA treatment, and binding of actin to several gene promoters is also detected in these conditions. Inhibitors targeting p38 mitogen-activated protein kinase prevent the increase in nuclear actin (Xu et al., 2010), but it is not known whether nuclear import or export of actin is affected.

Regulation of nuclear actin has also been linked to controlling transcription factor activity, and thereby to expression of specific sets of genes. Myocardin-related transcription factors (MRTFs) are co-activators of serum response factor (SRF), which regulate the expression of various cytoskeletal genes, including actin. MRTFs contain three RPEL repeats, which operate as G-actin sensors. In conditions where G-actin is abundant, actin monomer-binding to MRTF masks the nuclear localization signal (NLS) embedded within the RPEL domain, and thereby inhibits nuclear import of MRTF (Mouilleron et al., 2011; Pawlowski et al., 2010; Vartiainen et al., 2007). MRTF also interacts with actin inside the nucleus, where actin-binding promotes its nuclear export and prevents SRF activation (Vartiainen et al., 2007). Consequently, at high actin monomer concentrations, MRTF is mainly cytoplasmic and inactive. Reduction in actin monomer levels leads to nuclear localization of MRTF due to unmasking of its NLS and inhibited nuclear export, when actin-binding is prevented. In these conditions, MRTF can also activate SRF-mediated transcription (Vartiainen et al., 2007). Importantly, since actin itself, as well as many ABPs, are transcriptional targets of MRTF/SRF, this creates a feedback loop, where actin dynamics, in either cytoplasm or nucleus, controls the expression of proteins driving actin dynamics. Several mechanisms that culminate on nuclear actin have been reported to regulate MRTF/SRF-dependent gene expression. Serum stimulation causes transient polymerization of nuclear actin. This process is dependent on mDia formins, leads to MRTF nuclear accumulation and SRF activation (Baarlink et al., 2013). Also cell spreading induces mDia-dependent nuclear actin polymerization, and activates MRTF/SRF. Polymerization requires nuclear lamina components lamin A/C and emerin, as well as the LINC complex that connects the cytoskeleton to the nucleoskeleton, thereby linking nuclear F-actin formation to mechanosensing (Plessner et al., 2015). Another MRTF/SRF activating protein is MICAL-2, an atypical actin regulator that localizes to the nucleus and induces redox-dependent actin filament depolymerization, thus lowering nuclear G-actin levels (Lundquist et al., 2014). On the other hand, Ras association domain family 1 isoform A (RASSF1A) regulates nuclear actin levels, and thereby MRTF/SRF activity, by promoting the association between Exp6 and Ran GTPase. *RASSF1A* is silenced by promoter hypermethylation in many solid tumors, and cancer cells defective in this pathway display increased nuclear actin, reduced MRTF activity and consequent defects in cell adhesion (Chatzifrangkeskou et al., 2019). However, also activation of MRTF/SRF target genes have been reported in malignant cells to promote invasiveness. Nuclear capture of receptor tyrosine kinase EphA2-containing endosomes activates RhoG, which promotes phosphorylation of cofilin (Marco et al., 2021). Since unphosphorylated cofilin is required for nuclear import of actin (Dopie et al., 2012), this leads to decreased nuclear G-actin levels, and thereby activation of MRTF/SRF target gene expression (Marco et al., 2021). MRTF-A has been shown to promote vascular smooth muscle cell proliferation and migration (Smith et al., 2017). In these cells, elevated cAMP inhibits these processes by increasing the availability of cytoplasmic actin monomers, which are then transported into the nucleus, where they in turn inhibit YAP-TEAD and MRTF/SRF activity (McNeill et al., 2020). Thus, many mechanisms exist to regulate different aspects of nuclear actin from its levels to polymerization in order to control MRTF/SRF transcriptional activity.

Here we describe a novel cellular context, where nuclear actin levels are regulated to influence MRTF/SRF target gene expression. By using PC6.3 cells as an *in vitro* model for neuronal differentiation, we demonstrate that MRTF/SRF target gene expression is first rapidly activated, but then significantly repressed during the differentiation process. Repression of MRTF/SRF activity takes place at the level of MRTF-A subcellular localization, and coincides with increased nuclear actin levels due to decreased nuclear export of actin.

## Results

### Nuclear actin levels increase during PC6.3 cell differentiation

In this study, we use PC6.3 cells as an *in vitro* cell differentiation model. PC6.3 cells are a subclone of the rat PC12 pheochromocytoma cell line, and widely used to study the different pathways controlling neuronal differentiation and how factors, such as chemicals, affect neurite outgrowth. Treatment of PC6.3 cells with neural growth factor (NGF) ceases their proliferation, induces neurite outgrowth (Fig S1A) and the cells acquire properties of sympathetic neurons. Differentiation is induced through the TrkA receptor and initiates several signaling cascades (Zhou et al., 1995). Differentiation of PC6.3 cells can be easily followed by monitoring the number and length of neurites (Fig S1A), and by measuring the induction of several target genes (Dijkmans et al., 2008; Lee et al., 2005; see below Fig 2 for RNA-seq data). Differentiation is visible already 24 hours after NGF addition (Leopold et al., 2019) and reaches a plateau after 6 days (Pittman et al., 1993). Here we follow differentiation until 5 days after NGF treatment.

To study, if nuclear actin levels respond to PC6.3 differentiation we used cell fractionation (Fig 1A-B) and imaging approaches (Fig 1C-D). Cell fractionation revealed a significant increase in nuclear actin amounts after 3 days of differentiation. The levels were still elevated at 5 days of NGF treatment, but the variation was greater at later time points (Fig 1B). Similar observations were made by using fluorescence microscopy to quantify the distribution of GFP-actin between the nucleus and cytoplasm (Fig 1C-D). The ratio of nuclear to cytoplasmic GFP-actin intensity was the highest after 3 days of NGF treatment, and GFP-actin remains more nuclear at day 5 compared to untreated cells (Fig 1D). Western blotting of total actin levels revealed that the amount of actin decreases during PC6.3 cell differentiation and at day 4, the decrease is statistically significant (Fig S1B-C). This indicates that the increased nuclear actin is not due to overall increase in the amount of cellular actin, but rather reflects regulation of the subcellular distribution of actin. This idea is also supported by the results from GFP-actin localization (Fig 1C-D), which is not dependent on endogenous actin expression. Taken together, nuclear actin levels are subject to regulation during PC6.3 cell differentiation, reaching the peak level at 3 days after NGF addition.

**Figure 1.**
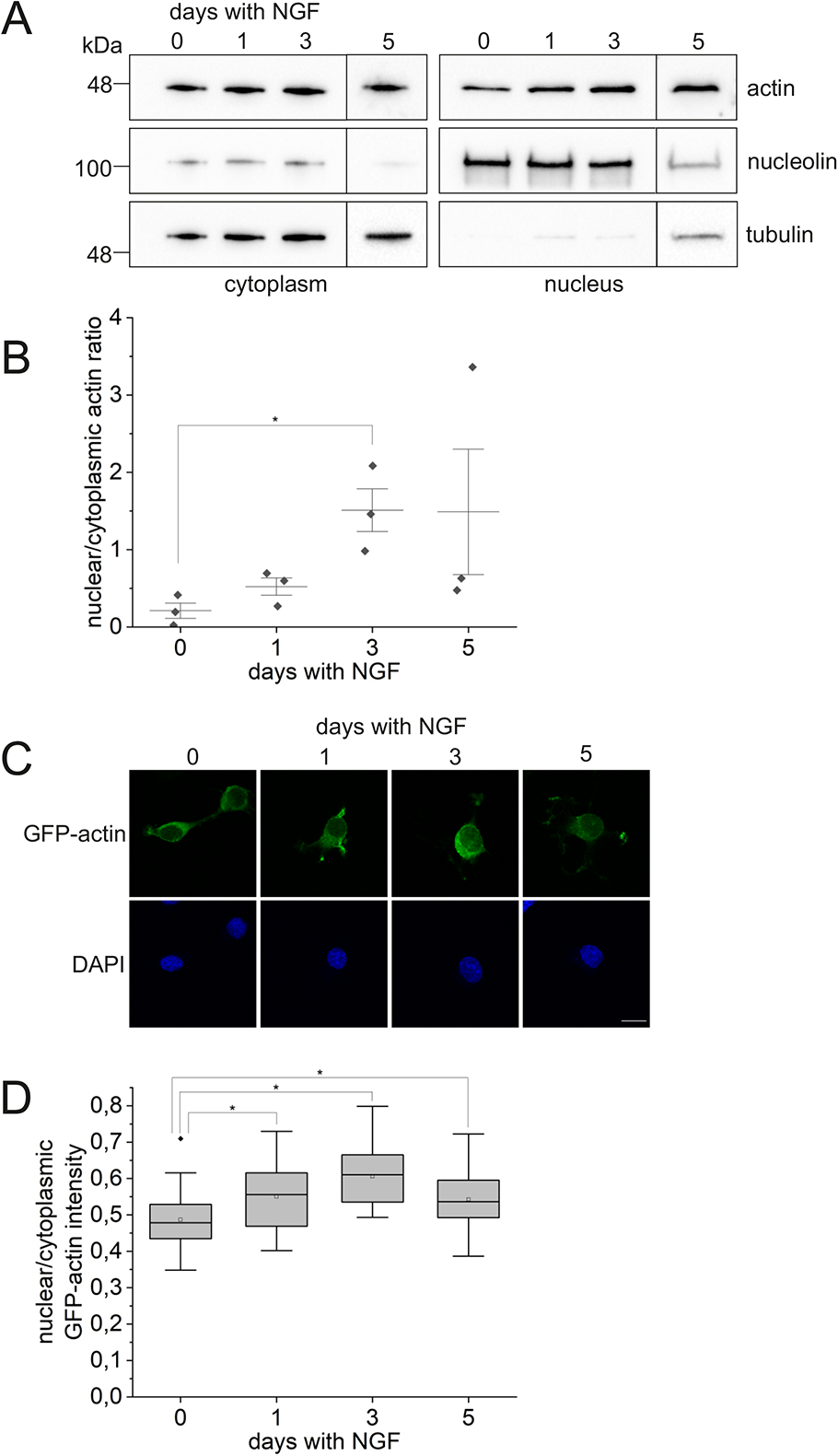
Nuclear actin levels increase during PC6.3 differentiation. A. Western blotting with indicated antibodies of nuclear and cytoplasmic protein fractions from PC6.3 cells differentiated for 0, 1, 3 or 5 days with NGF. Tubulin used as cytoplasmic and nucleolin as nuclear markers. Molecular weights based on markers are indicated on the left. Boxed areas are from separate Western blots. B. Quantification of nuclear to cytoplasmic actin distribution from Western blots. Data is shown as a scatter interval plot, with individual measurements (N=3) shown as black dots, error bars ± 0.5 standard deviation (SD) and mean as the horizontal line. * Statistically significant differences (P<0.05) with a two-sample t-test: 0 vs. 1 day P=0.146, 0 vs. 3 days P=0.042, 0 vs. 5 days P=0.305. C. Confocal microscopy images of PC6.3 cells transfected with GFP-actin and differentiated for 0, 1, 3 or 5 days. Nuclei stained with DAPI. Scale bar 10 µm. D. Quantification of GFP-actin distribution between nucleus and cytoplasm during PC6.3 cell differentiation. Data is shown as a box plot, where boxes represent 25%-75% of the values and error bars the range within 1.5IQR. Middle line is the median, open square is the mean, and black dots are outliers. N≥20 cells measured per condition. * Statistically significant differences (P<0.05) with a two-sample t-test: 0 vs. 1 day P=0.039, 0 vs. 3 days P=3.162E-5 and 0 vs. 5 days P=0.035. See also Fig S1.

### Reduced expression of MRTF/SRF target genes during PC6.3 cell differentiation

To explore the possible functional implications of increased nuclear actin levels during PC6.3 cell differentiation, we used RNA-sequencing (RNA-seq) to analyze changes in gene expression levels. RNA-seq was performed in four biological replicates from undifferentiated cells and cells treated with NGF for 1-5 days. Principal component analysis reveals that the largest differences in gene transcription take place during the first day of differentiation (Fig S2A). Analysis of the 12872 expressed genes (baseMean expression levels above 10) by comparing their expression to undifferentiated cells (log2FC) during NGF treatment (Supplementary Table S1) identifies different classes of transcriptional responses during PC6.3 cell differentiation (Fig S2B). These classes include, for example, genes that are either up- or downregulated already from the first day of differentiation (upper- and lower part, respectively, of Fig S2B), and those that show a more complex expression pattern, such as initial upregulation at day 1 of NGF treatment, followed by downregulated expression at subsequent days of differentiation (see below). In agreement with previous gene expression analysis of PC6.3 and the related PC12 cells (Dijkmans et al., 2008; Lee et al., 2005; Offermann et al., 2016), the genes upregulated already during the first day of NGF treatment (335 genes with logFC>1) contain many genes implicated in neuronal differentiation (for example GO terms nervous system development, P-value=0.0011; myelin sheath, P-value=0.0001; Supplementary table S2). Many of these genes, including *Syn2*, *Ngef, Nefm* and *Thy1*, remain upregulated until day 5 (209 from 335 with logFC>1 on day 1 and day 5). In addition to neuronal differentiation, the upregulated genes are also enriched for GO terms related to plasma membrane remodeling (Supplementary table S2), likely reflecting the need to remodel the cell surface for neurite extension. On the other hand, genes that display reduced expression already after 1 day of NGF treatment are linked to processes of cell division (for example GO terms cell division P-value=1.8E-13; chromosome segregation P-value=7.70E-12) in line with ceased proliferation during PC6.3 cell differentiation (Eggert et al., 2000). Overall, these results reflect the cellular processes known to take place during PC6.3 cell differentiation, and are in line with previous gene expression analysis in similar experimental systems (Dijkmans et al., 2008; Lee et al., 2005).

Among the genes that were downregulated at day 5 compared to day 1 of differentiation (Supplementary Table S2), we recognized several well-established target genes of MRTF/SRF, including for example *Actb*. Since nuclear actin has been shown to regulate MRTF/SRF transcriptional activity in several experimental systems, we decided to analyze expression of these genes in more detail. In the absence of an established target gene list for MRTF/SRF in neuronal cells, we utilized data from mouse fibroblasts, where MRTF target genes were defined based on binding of MRTF-A to these genes by ChIP-seq and their sensitivity to actin-binding drugs Cytochalasin D, which activates MRTFs and Latrunculin A, which inhibits MRTFs (Esnault et al., 2014). Of the 467 MRTF targets, 113 genes displayed at least 50% change in their expression during differentiation (Fig 2A-B; Supplementary table S3; based on the log2FC<-0.6 and log2FC>0.6 compared to non-treated cells). Of these genes, 46 were upregulated and 14 were downregulated at day 1 of NGF treatment. At day 3 of NGF treatment, there were 41 upregulated and 21 downregulated, and at day 5, 45 upregulated and 32 downregulated MRTF target genes, when compared to undifferentiated conditions. Thus, the number of downregulated MRTF target genes increased during PC6.3 cell differentiation. Moreover, majority (34 of 46) of the initially upregulated target genes displayed decreased expression, when comparing the day 5 to the day 1 of differentiation. This indicates that even though some MRTF/SRF target genes are initially activated during PC6.3 cell differentiation, there is an overall trend towards downregulating this pathway.

**Figure 2.**
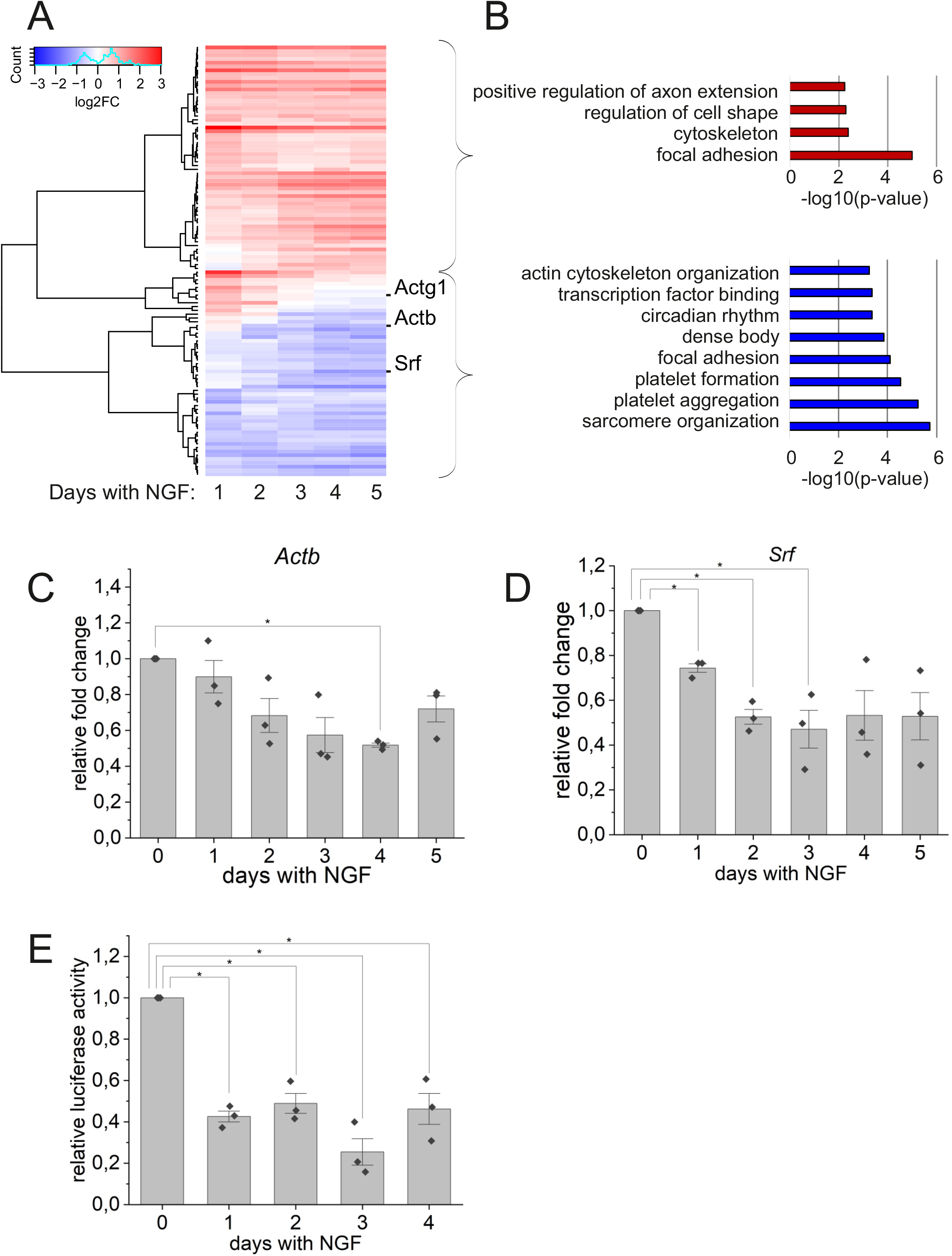
RNA-seq reveals decreased expression of MRTF/SRF target genes upon PC6.3 cell differentiation. A. Heatmap showing relative expression of MRTF-A direct target genes (Esnault et al. 2014), which display at least 50% change in their expression during differentiation (log2FC<-0.6 and log2FC>0.6; N=113) compared to undifferentiated cells. Canonical MRTF/SRF target genes indicated on the right. B. Gene Ontology enrichment analysis of MRTF-A direct target genes (see A) with increased (top) and decreased (bottom) relative expression during differentiation. C. qPCR of *Actb* during 0-5 days of differentiation normalized to undifferentiated control sample. Data is mean (N=3), error bars SD, individual measurements shown as black dots. * Statistically significant differences (P<0.05) with a one-sample t-test: 0 vs. 1 day P=0.436, 0 vs. 2 days P=0.101, 0 vs. 3 days P=0.063, 0 vs. 4 days P=7.827E-4, 0 vs. 5 days P=0.079. D. qPCR of *Srf* (N=3) during 0-5 days of differentiation normalized to undifferentiated control sample; data shown as in C. * Statistically significant differences (P<0.05) with a one-sample t-test: 0 vs. 1 day P=0.007, 0 vs. 2 days P=0.006, 0 vs. 3 days P=0.032, 0 vs. 4 days P=0.067, 0 vs. 5 days P=0.061. E. Relative SRF luciferase activity in PC6.3 cells differentiated for 0-4 days. Data is shown as mean (N=3), normalized to undifferentiated control sample, error bars SD and individual measurements shown as black dots. * Statistically significant differences (P<0.05) with a one-sample t-test: 0 vs. 1 day P=0.003, 0 vs. 2 days P=0.011, 0 vs. 3 days P=0.010, 0 vs. 4 days P=0.025. See also Fig S2.

To confirm decreased MRTF/SRF target gene expression, we performed qPCR analysis of selected target genes during 0-5 days of NGF treatment. In agreement with RNA-seq, the expression of *Actb* (Fig 2C) mRNA was decreasing during PC6.3 cell differentiation and is in line with total actin protein levels (Fig S1B-C). Also expression of *Srf* mRNA was downregulated from the first day of differentiation (Fig 2D). However, Western blotting of SRF protein levels did not reveal statistically significant changes (Fig S2C-D), although the trend was the same as observed for mRNA (Fig 2D). This indicates that the decreased expression of *Srf* mRNA is not sufficient to significantly alter SRF protein levels in this experimental set-up. Finally, we investigated the SRF transcriptional activity with a reporter assay in differentiating PC6.3 cells. In agreement with RNA-seq and qPCR data, SRF reporter activity declined already during the first day of differentiation, and the decrease continued towards day 4 of differentiation (Fig 2E).

Based on the RNA-seq data, some MRTF/SRF target genes were upregulated during the first day of differentiation, but then their expression declined from the second day of NGF treatment (Fig 2A-B). Due to this observation, and since previous studies have reported upregulation of SRF activity during the first hours of PC12 cell differentiation (Offermann et al., 2016), we decided to study also earlier time points during the first day of NGF treatment. Indeed, we observed increased SRF reporter gene activity after 8 hours of NGF stimulation compared to undifferentiated cells (Fig 3A). Moreover, qPCR analysis revealed that transcription of both *Srf* (Fig 3B) and *Vcl* (Fig 3C) were activated during the first hours of NGF stimulation. Especially *Srf* expression rapidly decreased back to baseline expression levels after peaking at 2 hours of NGF stimulation (Fig 3B), while *Vcl* expression remained elevated for longer time (Fig 3C). Taken together, our RNA-seq, qPCR and SRF reporter gene assays reveal that MRTF/SRF-mediated transcription is tightly regulated during PC6.3 cell differentiation. During the first hours of NGF-induced differentiation, expression of MRTF/SRF target genes is transiently activated, but then the pathway is repressed.

**Figure 3.**
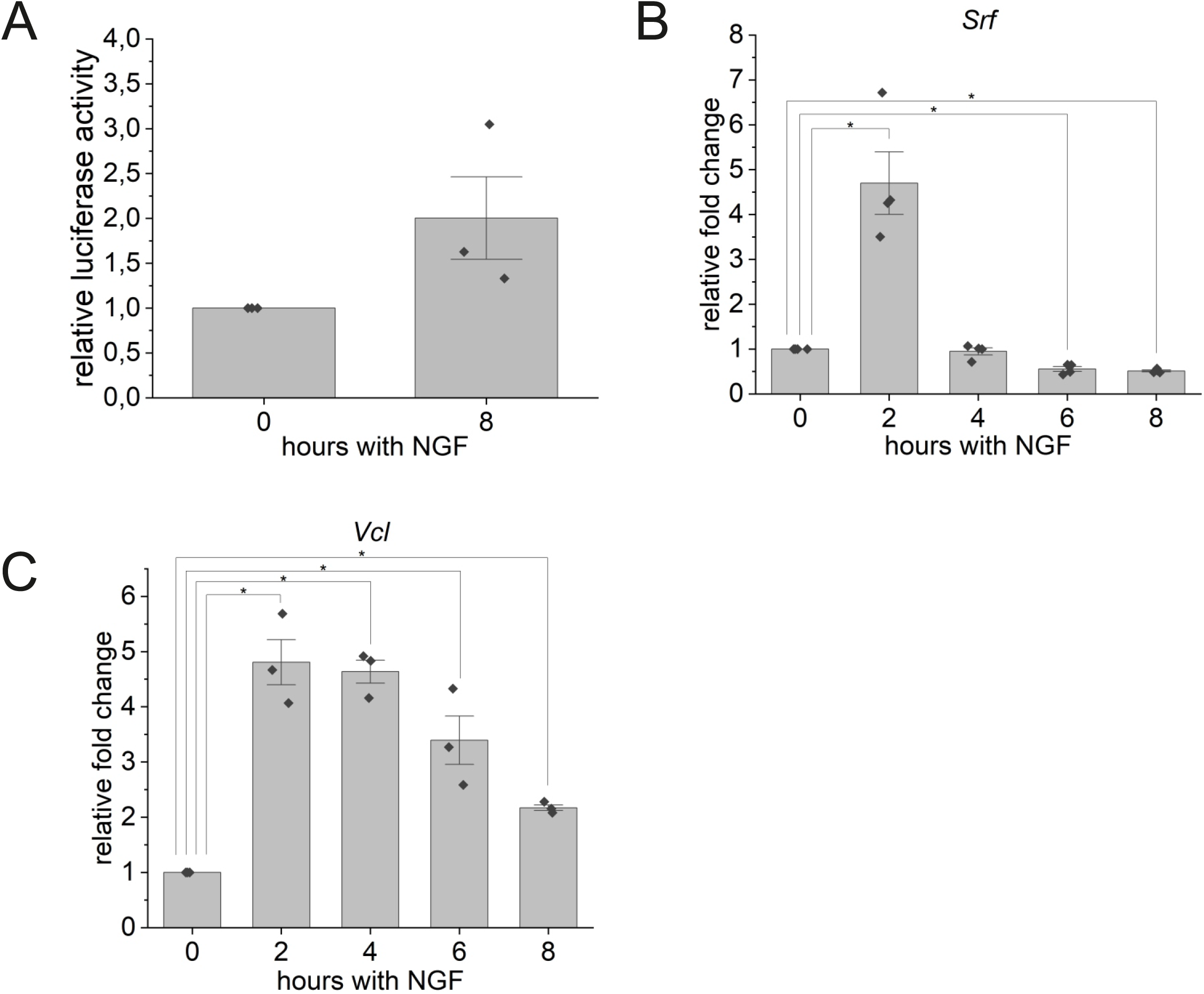
Increased MRTF/SRF transcriptional activity at early time points of NGF stimulation. A. Relative SRF luciferase activity in PC6.3 cells treated or not for 8 hours with NGF. Data is shown as mean (N=3), normalized to untreated sample, error bars SD and individual measurements shown as black dots. 0 vs. 8 hours P=0.199 with one-sample t-test. B. qPCR of *Srf* during 0-8 hours of differentiation. Data is shown as mean (N=4), normalized to undifferentiated control sample, error bars SD, individual measurements shown as black dots. * Statistical significance (P<0.05) with a one-sample t-test 0 vs. 2 hours P=0.013, 0 vs. 4 hours P=0.564, 0 vs. 6 hours P=0.004, 0 vs. 8 hours P=1.427E-4. C. qPCR of *Vcl*, shown as in B, (N=3). * Statistical significance (P<0.05) with a one-sample t-test: 0 vs. 2 hours P=0.015, 0 vs. 4 hours P=0.004, 0 vs. 6 hours P=0.042, 0 vs. 8 hours P=0.002.

### Subcellular localization of MRTF-A is subject to regulation during PC6.3 cell differentiation

Next, we wanted to understand the cellular processes that lead to regulation of MRTF/SRF target genes during PC6.3 cell differentiation. Since SRF protein levels were not significantly reduced during the process (Fig S2C-D), it is unlikely to fully explain reduced expression of its target genes. We therefore turned our attention to MRTF cofactors. RNA-seq did not reveal significant changes in *Mrtf-a* expression (Supplementary Table S1), and also Western blotting with antibody recognizing MRTF-A failed to reveal consistent changes in protein levels during 0-5 days of PC6.3 cell differentiation, apart from a transient increase in MRTF-A expression at day 1 (Fig 4A-B). It is well established in fibroblasts that nuclear actin regulates subcellular localization of MRTF-A (Baarlink et al., 2013; Vartiainen et al., 2007), but the nucleo-cytoplasmic shuttling process of MRTF in neuronal cells seems to be cell-type specific (Knoll, 2010). We therefore used immunofluorescence staining of MRTF-A to study its subcellular localization during PC6.3 cell differentiation. In agreement with target gene expression (Fig 2A, C-E and Fig 3), MRTF-A was translocated from the cytoplasm to the nucleus already after 30 min of NGF treatment, but then its localization returned to almost baseline at 2 hours of NGF (Fig 4C-D). Experiments with longer time points revealed reduced nuclear MRTF-A compared to untreated cells from day 3 of differentiation (Fig 4E-F). Taken together, MRTF-A subcellular location is tightly regulated during PC6.3 cell differentiation. During the first hours of differentiation, MRTF-A transiently accumulates in the nucleus and then returns back to baseline. Around day 3 of differentiation, the nuclear levels of MRTF-A are even further reduced. This localization data fits very well with MRTF/SRF target gene expression, and repression of the pathway coincides with increased nuclear actin levels, in agreement with data from other cellular systems with elevated nuclear actin (Chatzifrangkeskou et al., 2019; McNeill et al., 2020).

**Figure 4.**
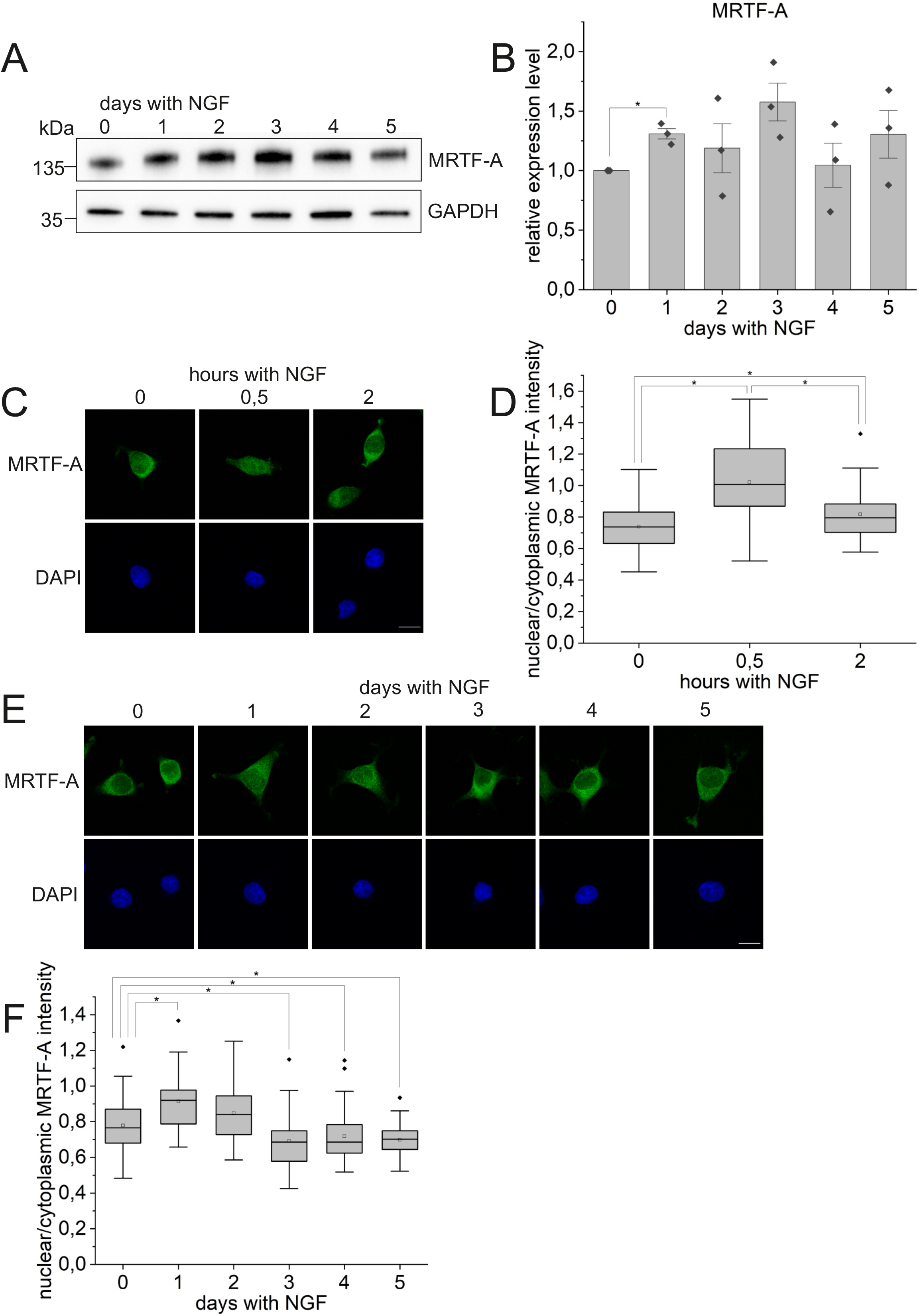
MRTF-A nuclear localization is regulated during PC6.3 cell differentiation. A. Western blotting of total MRTF-A protein levels during PC6.3 cell differentiation at indicated days with NGF. GAPDH used as the loading control. Note that GAPDH is the same as in Fig S2C. B. Quantification of MRTF-A levels from Western blots. Data is mean (N=3), normalized to undifferentiated sample, error bars SD, individual data points shown as black dots. * Statistically significant differences (P<0.05) with a one-sample t-test. 0 vs. 1 day P=0.025, 0 vs. 2 days P=0.510, 0 vs. 3 days P=0.088, 0 vs. 4 days P=0.854, 0 vs. 5 days P=0.320. C. Confocal microscopy images of cells differentiated for 0-2 hours and stained with anti-MRTF-A antibody. Nuclei stained with DAPI. Scale bar 10 µm. D. Quantification of MRTF-A nuclear to cytoplasmic distribution from microscopy images. Data is shown as a box plot, where boxes represent 25%-75%, middle line is the median, open square is the mean, black dots are the outliers and error bars show the range within 1.5IQR. N≥36 cells measured per condition. * Statistically significant differences (P<0.05) with a Mann-Whitney test: 0 vs. 0.5 h P=1.687E-7, 0 vs. 2 h P=0.011, 0.5 vs. 2 h P=1.613E-4. E. Confocal microscopy images of cells differentiated for 0-5 days and stained with anti-MRTF-A and DAPI. Scale bar 10 µm. F. Quantification of MRTF-A nuclear to cytoplasmic distribution. Data is shown as in D; N≥30 cells measured per condition. * Statistically significant differences (P<0.05) with a Mann-Whitney test: 0 vs. 1 day P=0.001, 0 vs. 2 days P=0.065, 0 vs. 3 days P=0.012, 0 vs. 4 days P=0.046, 0 vs. 5 days P=0.021.

### Reduced nuclear export of actin upon PC6.3 cell differentiation

We have previously shown that nuclear actin levels are actively maintained by nuclear transport (Dopie et al., 2012). Thus, the increase in nuclear actin levels observed during PC6.3 cell differentiation can be due to either increased nuclear import or decreased nuclear export of actin. To study this, we used different photobleaching methods to measure nuclear transport of GFP-actin (Skarp & Vartiainen, 2013). These assays were performed until day 3 of differentiation, when the actin accumulation in the nucleus seems to peak. First, we studied nuclear import of actin with fluorescence recovery after photobleaching (FRAP) assay. Here, the nucleus is bleached once with full laser power, and the recovery of nuclear fluorescence, due to import of unbleached GFP-actin molecules, is followed. As described before (Dopie et al., 2012), the beginning of the recovery curve gives a good measure of the nuclear import rate, but later nuclear export will start to influence fluorescence recovery. Overall, the shape of the recovery curve was very similar in cells differentiated for 0, 1 and 3 days (Fig 5A), and also the import rate between different days did not show significant changes (Fig 5B). We note that the average fluorescence recovery curve for the cells treated for 1 day with NGF shows somewhat higher initial fluorescence levels, which may reflect subtle differences in overall actin mobility in this sample.

**Figure 5.**
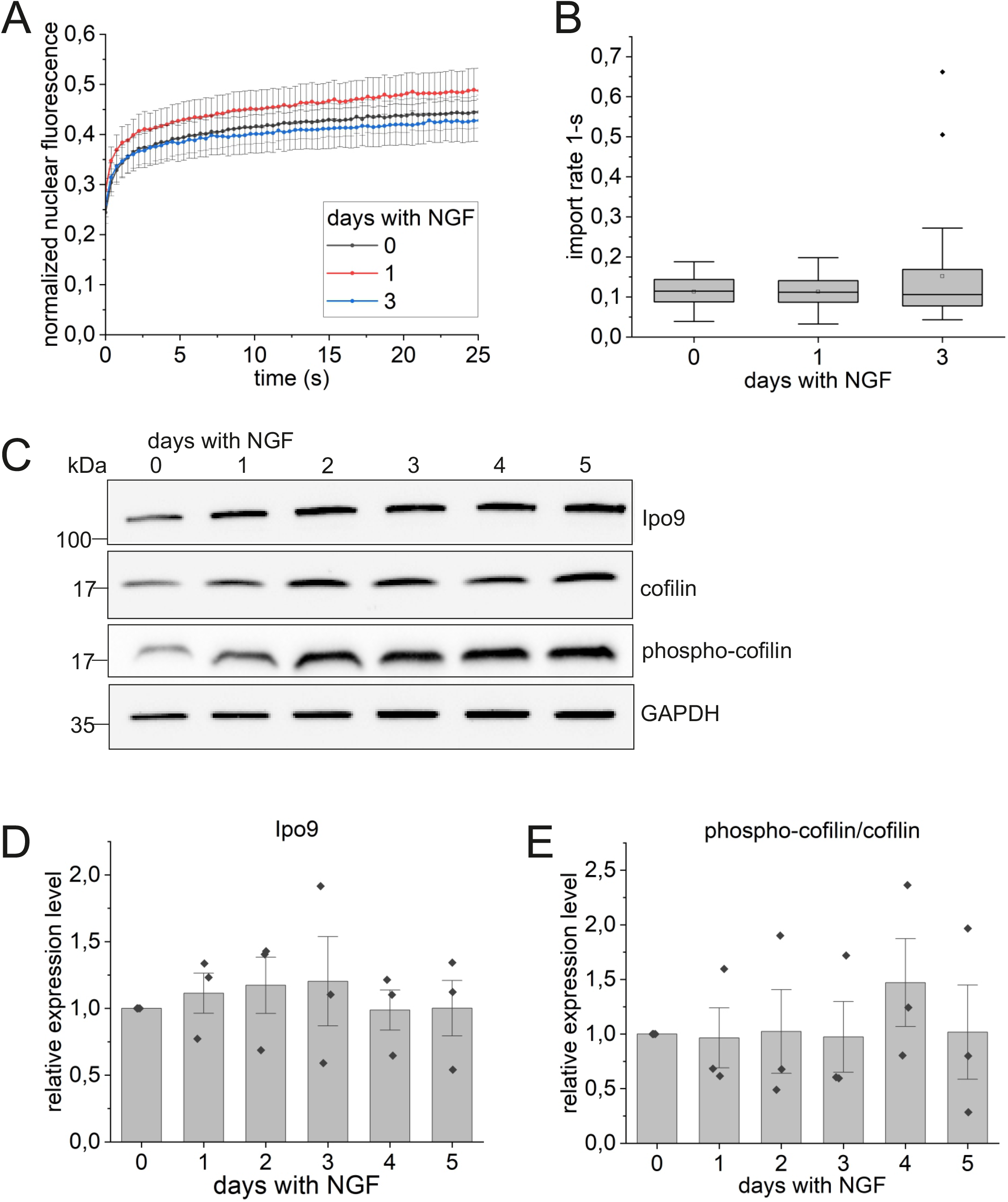
Nuclear import of actin is not affected during PC6.3 cell differentiation. A. Fluorescence recovery curves from FRAP assay representing nuclear import of fluorescent actin molecules in cells differentiated for indicated days with NGF. Data is mean (N≥27 cells measured per condition), normalized to pre-bleach values, error bars SD. B. Apparent nuclear import rate of GFP-actin from FRAP assay in cells differentiated for indicated days with NGF. Data is shown as a box plot, where boxes represent 25%-75%, middle line is the median, open square the mean, error bars represent the range within 1.5IQR and black dots are the outliers. Statistical significance with a Mann-Whitney test: 0 vs. 1 day P=0.885, 0 vs. 3 days P=0.997. C. Western blot of Ipo9, cofilin and phospho-cofilin levels during PC6.3 differentiation for indicated days. GAPDH used as the loading control. All proteins blotted from the same sample set. Molecular weights on the left. D. Quantification of Ipo9 Western blots. Data is mean (N=3), normalized to undifferentiated sample, error bars SD and individual data points shown as black dots. Statistical significance with a one-sample t-test 0 vs. 1 day P=0.578, 0 vs. 2 days P=0.550, 0 vs. 3 days P=0.651, 0 vs. 4 days P=0.950, 0 vs. 5 days P=0.994. E. Quantification of phospho-cofilin/cofilin levels from Western blots. Data (N=3) is shown as in D. Statistical significance with a one-sample t-test: 0 vs. 1 day P=0.923, 0 vs. 2 days P=0.963, 0 vs. 3 days P=0.950, 0 vs. 4 days P=0.418, 0 vs. 5 days P=0.976. See also Fig S5.

Our previous research has demonstrated that nuclear import of actin is dependent on Ipo9 and cofilin, in its unphosphorylated form (Dopie et al., 2012). To study the abundance of these molecules during PC6.3 cell differentiation, we used Western blotting (Fig 5C). However, we did not observe any significant changes in either Ipo9 protein levels (Fig 5D), or the ratio between phospho-cofilin and cofilin (Fig 5E). The total levels of cofilin are slightly higher during the first days of differentiation (Fig S5A), but these changes are not statistically significant. Furthermore, this applies also to phospho-cofilin (Fig S5B). Moreover, based on RNA-seq data (Supplementary table S1) neither *Ipo9* nor *Cfl* mRNAs displayed significant changes in their expression levels during PC6.3 cell differentiation. This was further confirmed for *Ipo9* using qPCR (Fig S5C). Combined, these experiments did not show any significant changes in nuclear import rate of actin (Fig 5B) or in the abundance of proteins required for this process (Fig 5D-E). This suggests that nuclear import of actin is not the regulatory step driving increased nuclear actin levels during PC6.3 cell differentiation.

Next, we studied nuclear export of actin with fluorescence loss in photobleaching (FLIP) assay, where cytoplasm is continuously bleached, and loss of nuclear fluorescence, due to nuclear export of GFP-actin, is measured (Dopie et al., 2012; Skarp & Vartiainen, 2013). This assay revealed clear differences especially at 3 days of NGF treatment compared to untreated cells (Fig 6A-C). Nuclear export rate, plotted from the initial loss of fluorescence, was significantly slower at 3 days of NGF treatment compared to untreated cells (Fig 6B). We also note that the export rates were clearly less variable in differentiated cells compared to undifferentiated. Moreover, there was significantly more GFP-actin fluorescence remaining in the nucleus after 100 s of bleaching already in the cells treated for 1 day with NGF, and the effect was even more pronounced at day 3 (Fig 6C). In addition to the nuclear export rate, the portion of GFP-actin remaining in the nucleus at the end of the assay represents the availability of export competent actin monomers (Dopie et al., 2012). Our results therefore suggest that in differentiated cells, actin might be more polymerized or tightly bound to nuclear complexes, such as chromatin remodelers, than in undifferentiated cells. Since our data clearly indicates decreased nuclear export of actin in differentiated PC6.3 cells compared to undifferentiated cells, studies on Exp6 protein levels would have been very interesting. However, we failed to identify an antibody that would have specifically detected Exp6 protein from the rat cells used for this study. Due to the strict cut-off *Xpo6* was not among the differentially expressed genes in the RNA-seq dataset (Supplementary table S1). Nevertheless, qPCR analysis revealed moderate decrease in *Xpo6* mRNA levels during PC6.3 cell differentiation (Fig 6D), which is in line with the RNA-seq data (for example at day 3, log2FC=-0.35 with P-value = 0.0000002 compared to undifferentiated cells). Nevertheless, whether this decrease is sufficient to impact Exp6 protein levels, and thereby nuclear export of actin, warrants the development of new reagents.

**Figure 6.**
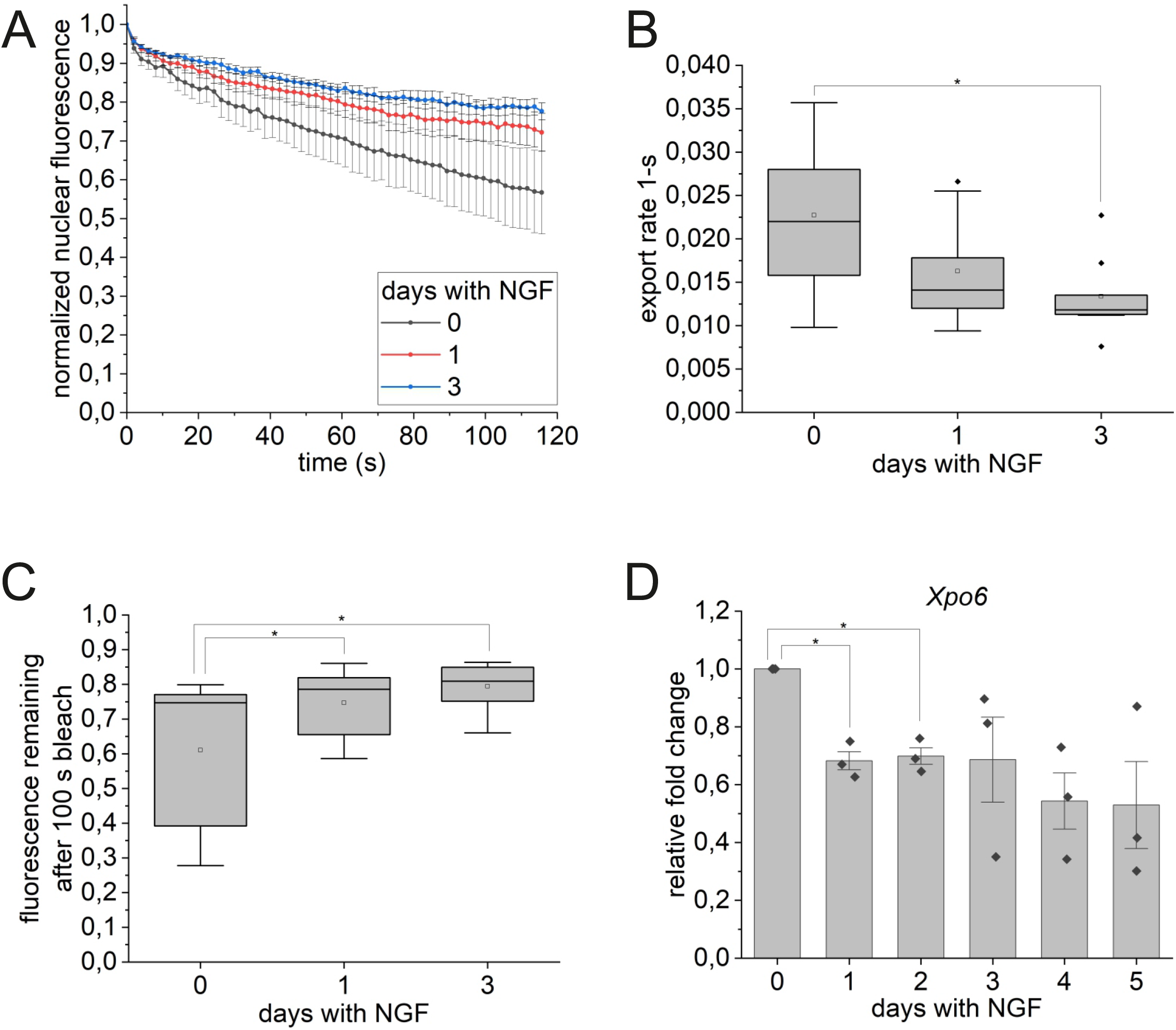
Slower nuclear export of actin during PC6.3 cell differentiation. A. Fluorescence loss curves from FLIP assay representing nuclear export of actin molecules in cells differentiated for indicated days with NGF. Data is mean (N≥11 cells per condition), normalized to pre-bleach frames and error bars SD. B. Nuclear export rate of GFP-actin quantified from FLIP assay. Data is shown as a box plot, where boxes represent 25%-75%, middle line is the median, open square the mean, error bars represent the range within 1.5IQR and black dots are the outliers. * Statistically significant differences (P<0.05) with a two-sample t-test: 0 vs. 1 day P=0.092, 0 vs. 3 days P=0.014. C. Fluorescence remaining after 100 s bleach in the FLIP assay. Data is shown as a box plot as in B. * Statistically significant differences (P<0.05) with a Mann-Whitney test: 0 vs. 1 day P=0.027, 0 vs. 3 days P=0.003. D. qPCR of *Xpo6* mRNA (N=3) during PC6.3 cell differentiation. Data is shown as mean (N=3), normalized to undifferentiated control sample, error bars SD and individual data points shown as black dots. * Statistically significant differences (P<0.05) with a one-sample t-test: 0 vs. 1 day P=0.013, 0 vs. 2 days P=0.012, 0 vs. 3 days P=0.206, 0 vs. 4 days P=0.055, 0 vs. 5 days P=0.113.

## Discussion

The cellular actin balance is maintained by active nucleo-cytoplasmic shuttling that is subject to regulation at different levels to control the relative abundance of actin in the nuclear and cytoplasmic compartments. Indeed, regulation of nuclear actin levels have been reported during various cellular processes, from differentiation to disease (Hyrskyluoto & Vartiainen, 2020). Functionally, alterations in nuclear actin levels are most often linked to either gene-specific transcriptional regulation or overall transcriptional activity. Nevertheless, neither the signaling pathways nor the mechanisms by which increased or decreased nuclear actin levels influence transcription are fully understood. Here we extend the studies on nuclear actin regulation by demonstrating that nuclear actin levels increase upon PC6.3 cell differentiation towards neuron-like cells (Fig 1). This increase is due to reduced nuclear export of actin during differentiation (Fig 6). Gene expression analysis reveals that MRTF/SRF target gene expression is first transiently activated (Fig 3), but then substantially downregulated during PC6.3 cell differentiation (Fig 2). This study therefore provides novel insights into nuclear actin regulation and MRTF/SRF pathway in a cellular differentiation model.

The nucleo-cytoplasmic shuttling process of actin offers several regulatory points to influence the cellular actin balance (Dopie et al., 2012). Indeed, previous studies have revealed regulation at the level of both nuclear import (Le et al., 2016) and export (Bohnsack et al., 2006; Fiore et al., 2017), and through regulation of either nuclear transport factors (Fiore et al., 2017) or actin itself (Le et al., 2016). By using photobleaching assays to measure nuclear transport rates of actin, we show that in PC6.3 cells, actin is regulated at the level of nuclear export (Fig 6A-C), but not nuclear import (Fig 5A-B). Reduced export could be achieved by modulating either the levels and/or activity status of the proteins needed for nuclear export of actin, or by regulating the availability of transport competent actin. Our fluorescence loss in photobleaching experiments reveal both decreased export rate of actin (Fig 6B) and increased retention of actin in nucleus (Fig 6C). These two processes are obviously linked, but the FLIP assay used here does not allow us to differentiate between them. In *Xenopus* oocytes, the massive nuclear actin amount that gives rise to a filamentous meshwork is achieved by post-transcriptional downregulation of Exp6 protein levels (Bohnsack et al., 2006). Our qPCR results suggest a modest decrease in *Xpo6* mRNA levels during PC6.3 cell differentiation (Fig 6D). However, our failure to identify an antibody that would reliably recognize rat Exp6 protein prevented further studies on this topic. RASSF1A is required for nuclear export of actin by supporting the interaction between Exp6 and the Ran GTPase, and its expression levels are regulated by promoter hypermethylation in several solid tumors (Chatzifrangkeskou et al., 2019). However, our RNA-seq analysis did not reveal significant changes in *Rassf1* during PC6.3 cell differentiation (Supplementary table S1). In mouse epithelial cells, attenuation of the PI 3-kinase activity enhances nuclear export of actin (Fiore et al., 2017), but the mechanism has remained unclear. Interestingly, NGF seems to stimulate PI3K activity during PC6 and PC12 cell differentiation (Nusser et al., 2002; Yan et al., 2020), and the PI3K inhibitor LY294002 inhibits NGF-induced neurite outgrowth (Shi & Andres, 2005). It is therefore tempting to speculate that enhanced PI3K activity during PC6.3 differentiation reduces nuclear export actin and leads to the observed accumulation of nuclear actin. Another signaling pathway that could play a role is the MAP kinase pathway, which is also activated downstream of NGF (Xing et al., 1998). Upon differentiation of HL-60 cells towards macrophages, treatment with MAP kinase inhibitors prevents nuclear accumulation of actin (Xu et al., 2010). Development of new tools to study the Exp6 protein is required to explore these possibilities further.

Actin monomer levels can limit nuclear transport rates of actin (Dopie et al., 2012) and our FLIP assay hinted that less actin might be available for nuclear export in differentiated PC6.3 cells (Fig 6C). This could arise either from increased nuclear polymerization of actin or from actin-binding to nuclear complexes. We did not observe clear phalloidin staining in the nuclei of PC6.3 cells in any condition (data not shown). Moreover, nuclear actin polymerization is most often linked to activation of MRTF/SRF target gene expression (Baarlink et al., 2013; Plessner et al., 2015), which is contrary to what we observe here in PC6.3 cells (Fig 2A, C-D). Since actin is an essential component of chromatin remodelers (Klages-Mundt et al., 2018) and also needed to sustain maximal transcription (Dopie et al., 2012; Sokolova et al., 2018) it can be speculated that these roles become more important during differentiation, requiring more actin to be associated with these complexes. This could explain increased retention of actin during PC6.3 cell differentiation. It is indeed known that during differentiation, thousands of genes are repositioned inside the nucleus and these relocations are driven by chromatin remodeling and correlate with changes in transcription and replication timing (Therizols et al., 2014). Actin could be one possible candidate to regulate these global changes in chromatin organization during differentiation. Indeed, mouse embryonic fibroblasts (MEFs) lacking the β-actin gene display altered 3D genome architecture (Mahmood et al., 2021) and are defective in their reprogramming capacity towards osteoblast-like cells (Gjorgjieva et al., 2020), adipocytes (Al-Sayegh et al., 2020) and chemical-induced neurons. In this latter case, β-actin deficiency is linked to loss of chromatin-binding of Brg1, an ATPase of the BAF chromatin remodeling complex. This leads to defects in heterochromatin marks and impaired expression of neuronal gene programs (Xie et al., 2018).

Whether PC6.3 cells undergo profound changes in their global chromatin organization is not known at the moment, but our RNA-seq analysis revealed 1338 genes that displayed altered expression (Log2FC>1 or <-1) during PC6.3 cell differentiation. Functional annotation of these genes reflects the cellular processes that are known to take place during differentiation: decreased proliferation and increased plasma membrane remodeling for neurite outgrowth (Fig 2B). Whether increased nuclear actin is required to elicit these changes in gene expression is not known. In several other experimental systems, modulation of nuclear actin levels have been linked to regulation of MRTF/SRF target gene expression (Chatzifrangkeskou et al., 2019; Lundquist et al., 2014; Marco et al., 2021). Hence it was not too surprising that we observed down-regulation of MRTF/SRF transcription activity (Fig 2E) and target gene expression (Fig 2A, C-D) during later stages of PC6.3 cell differentiation, when increased nuclear actin was also observed. Mechanistically, this down-regulation of MRTF/SRF is achieved by controlling MRTF-A nuclear localization, which was markedly decreased during differentiation (Fig 4E-F). Interestingly, regulation of MRTF subcellular localization in neuronal cells seems to be more complex and cell type-specific than in fibroblasts, where this process has been extensively studied. In some neuronal cells, MRTF-A has been shown to shuttle between the cytoplasm and the nucleus, whereas in cortical and hippocampal neurons, exclusively nuclear localization has been reported (Knoll, 2010). It has been hypothesized that due to the mostly nuclear MRTF-A localization in some cells, nuclear actin would modulate MRTF-A expression rather than localization (Stern et al., 2009). However, in our PC6.3 cell model, MRTF-A protein levels do not change upon differentiation (Fig 4B), and MRTF-A shows both cytoplasmic and nuclear distributions (Fig 4C, E), which seem similar to the actin-dependent nucleo-cytoplasmic shuttling characterized in fibroblasts (Vartiainen et al., 2007).

Importantly, before the decreased activity of MRTF/SRF after days with NGF, the pathway is transiently activated during the first hours of applying the differentiation conditions (Fig 3). Activation of MRTF/SRF downstream of NGF has been shown already before (Lundquist et al., 2014; Wickramasinghe et al., 2008), and several studies have reported the requirement for MRTF/SRF target gene expression during neuronal differentiation and function. For example, expression of dominant-negative MRTF constructs inhibits neuronal outgrowth and guidance (Knoll et al., 2006) and depletion of MRTFs by RNAi reduces dendritic complexity (Ishikawa et al., 2010). Moreover, lack of both MRTFs in mice leads to disruption of multiple brain areas with defects in neuronal migration and neurite outgrowth (Mokalled et al., 2010). In addition, expression of actin mutants in either cytoplasm or nucleus regulates neuronal motility in SRF-dependent manner (Stern et al., 2009). Since MRTF/SRF regulate the expression of numerous cytoskeletal target genes, defects in cellular processes that are driven especially by actin dynamics are therefore not surprising, and early activation of MRTF/SRF during neuronal differentiation is logical. It is, however, less obvious why MRTF/SRF activity decreases later during differentiation. It might be that this is merely a specific feature of the used cell model, and reflects the conditions used to stimulate differentiation by low serum containing media. On the other hand, a recent study utilizing reprogramming of somatic cells to induced pluripotent stem cells has suggested that SRF can repress expression of cell-type specific genes through their epigenetic regulation (Ikeda et al., 2018). Perhaps SRF inactivation is required also in the PC6.3 cells to allow expression of genes required for neuronal differentiation. An important caveat of our RNA-seq analysis is that MRTF/SRF target genes have not been firmly established in neurons. We used a target gene list from fibroblasts derived from MRTF-A chromatin-binding patterns and sensitivity to actin-binding drugs (Esnault et al., 2014). Target genes are often cell-type specific, due to, for example, partnership with different transcription factors. Overall, further studies are needed to establish MRTF/SRF target genes in different cellular contexts.

## Acknowledgements

We thank Paula Maanselkä for excellent technical assistance. This work was supported by Sigrid Juselius foundation, Jane and Aatos Erkko foundation and Academy of Finland grants 338281and 330254 to MKV, as well as Academy of Finland grant 308890 to AH and Instrumentarium Science foundation grant to SK. Imaging was performed at the Light Microscopy Unit, Institute of Biotechnology and RNA-seq at Biomedicum Functional Genomics Unit, both supported by HiLIFE and Biocenter Finland.

## Materials and methods

### Plasmids

GFP-actin plasmid is described in Dopie et al. 2012.

### Antibodies

The following antibodies were from Thermo Scientific: GAPDH (MA5-15738, 1:2500), the following ones from Merck: Anti-Actin AC-40 (A3853, 1:1000), tubulin (T6074, 1:2500) and nucleolin (N2662, 1:1000). SRF (5147, 1:750), MRTF-A (14760S, 1:1000) and cofilin (3312, 1:1000) antibodies were from Cell Signaling Technology, Ipo9 (PAB0154, 1:1000) from Abnova and MRTF-A G8 (sc-390324, 1:50) from Santa Cruz Biotechnology. Phospho-Ser3 cofilin antibody (11139-1, 1:1000) was from Signalway antibody. The following secondary antibodies were from Thermo Fisher Scientific: horseradish peroxidase (HRP) conjugated anti-mouse IgG (G-21040, 1:5000), HRP-conjugated anti-rabbit IgG (G-21234, 1:5000) and Alexa-Fluor-488 conjugated anti-mouse IgG (A11001, 1:500).

### Cell lines, differentiating cells and transfections

PC6.3 cells obtained from Urmas Arumäe (Tallinn University of Technology) were cultured in RPMI-1640 medium (Lonza) supplemented with 10% horse serum (HS; Gibco), 5 % fetal bovine serum (FBS; Gibco), penicillin-streptomycin (Thermo Fisher Scientific) and GlutaMAX (Thermo Fisher Scientific) in 5% CO_2_ incubator at + 37 °C. For differentiation, cells were plated on poly-L-lysine (Merck) and laminin (Merck) coated cell culture plates in full RPMI medium. Next day, medium was replaced to low serum RPMI (RPMI-1640 supplemented with 1% HS and 1% FBS) with 50 ng/ml NGF 2.5S Native Mouse Protein (Gibco) to differentiate cells for desired time. Control cells were kept in low serum RPMI without NGF during the duration of the experiment. For GFP-actin transfection, differentiating PC6.3 cells were transfected using JetPrime transfection reagent (Polyplus) according to manufacturer’s protocol using total DNA amount of 250 ng per 24 well plate well/1000 ng per 35 mm dish out of which 25% was GFP-actin and the rest filled with pEF-Flag plasmid.

### Nuclear/cytosol fractionation

PC6.3 cells were plated to 10 cm cell culture plates at densities 500 000 - 1 000 000 cells per plate, differentiated for 0-5 days and nuclei were separated from the cytoplasm using Nuclear/Cytosol Fractionation Kit (BioVision) according to manufacturer’s instructions. SDS-PAGE loading buffer was added to resulting fractions and the samples were processed for SDS-PAGE and Western blot using anti-Actin Ac-40, anti-tubulin and anti-nucleolin antibodies to verify the performance of the fractionation and to study actin distribution during differentiation. Intensities of bands were measured with Fiji ImageJ software, actin in cytoplasmic and nuclear fractions normalized to tubulin and nucleolin, respectively, and data presented as nuclear to cytoplasmic actin ratio.

### Immunofluorescence and microscopy

PC6.3 cells were plated on 24 well cell culture plates with coverslips at a density of 5 000-10 000 cells per well. Cells were differentiated for 0-8 hours or for 0-5 days for MRTF-A staining and for 0-5 days for GFP-actin transfection. The transfection was done one day before fixation. Before fixation, relief phase images were taken with FLoid Cell Imaging Station (Thermo Scientific) using a fixed 20x plan fluorite objective to confirm that cells were differentiated. Cells were fixed with 4% paraformaldehyde for 20 min, washed three times with PBS and permeabilized with 0.1% Triton X-100 (Merck) in PBS for 5 min. Samples were then blocked in blocking buffer [1% gelatin (Merck), 1% BSA (Merck), 10% FBS (Lonza) in PBS] for 30 min and mounted in Prolong Diamond with DAPI (Thermo Scientific), or stained with primary antibody MRTF-A G8 diluted in blocking buffer for 45 min, washed and stained with secondary Alexa Fluor-488 antibody for 45 min. Coverslips were washed in PBS and in MilliQ and mounted as above.

Imaging was done using Zeiss LSM 700 confocal with Axio Imager M2 microscope, ZEN 2012 software and LCI Plan-Neofluar 63x/1.30 Imm Corr glycerol objective. Fixed samples were imaged to analyze MRTF-A or GFP-actin distribution in cells. Imaging was done using 405 nm (for DAPI) and 488 nm (for Alexa Fluor-488 or GFP) lasers, the pinhole was set to 1 AU, resolution was 668×668 and bit depth 8. Fluorescence intensities were measured from cell nucleus and cytoplasm using Fiji ImageJ software and the ratios were calculated by dividing the average nuclear intensity with cytoplasmic intensity.

### Total protein amounts during differentiation

PC6.3 cells were plated on 6 well cell culture plates at a density of 80 000 cells per well and differentiated with NGF for 0-5 days. Differentiated cells were lysed to 1x SDS-PAGE loading buffer and samples were processed for SDS-PAGE and Western blotting using anti-Actin AC-40, MRTF-A (Cell Signaling Technology), SRF, Ipo9, cofilin, phospho-cofilin and GAPDH antibodies. Intensities of bands were measured with Fiji ImageJ software and normalized to GAPDH.

### Luciferase assay

Cells were plated on 24 well cell culture plates at a density of 100 000 cells per well and differentiated with NGF for 0-8 hours or 0-4 days. Cells were transfected with SRF reporter p3DA.luc (8 ng per well) and reference reporter pTK-RL (20 ng per well) by JetPrime transfection reagent. Total DNA amount was 200 ng per well, and the rest was filled with pEF-myc plasmid. For 0 and 8 h differentiated cells, the transfection was done the day before NGF stimulation. For 0-4 days differentiated cells, the transfection was done one day before the assay. After differentiation was completed, Dual-Luciferase reporter assay system (Promega) kit was used according to manufacturer’s instructions to measure the relative SRF reporter activity. For data analysis, the activity of firefly luciferase was normalized to renilla luciferase activity.

### FLIP

PC6.3 cells were plated on 35 mm cell culture dishes at a density of 80 000 cells per dish and differentiated for 0-3 days as previously. During the last day before imaging, cells were transfected with GFP-actin. Next day, cells were live imaged in + 37 °C, 5% CO_2_ in Okolab bold line cage incubator using Zeiss LSM 700 confocal with Axio Imager M2 microscope with W Plan-Apochromat 63x/1.0 dipping objective. Acquisition software was ZEN 2012. Imaging parameters: pinhole 1 AU, resolution 256×256, bit depth 12, speed 7, line average 1, zoom 2. The cytoplasm was continuously bleached for 120 s with 2 s intervals using 100% laser power (488 nm/10 mW) and the loss of fluorescence in the nucleus resulting from export of GFP-actin to the cytoplasm was recorded. Bleaching was started after 3 scans. Data was processed by setting the pre-bleach values to 1 and by producing a linear fit of the first data points to get the export rate.

### FRAP

PC6.3 cells were plated on 35 mm glass bottom cell culture dishes at a density of 80 000 cells per dish and differentiated for 0-3 days as previously. During the last day before imaging, cells were transfected with GFP-actin. Next day, cells were live imaged in + 37 °C, 5% CO_2_ in LifeImagingSystems incubation system using Leica TCS SP5 II HCS-A confocal with DMI6000 B microscope with HCX PL APO 63x/1,2 W Corr/0,17 CS water objective, RSP 500 beam splitter and FRAPbooster. Acquisition software was LAS AF 2.7.7. Imaging parameters: pinhole 1 AU, resolution 256×256, bit depth 12, speed 700 Hz, bidirectional X scan, line average 2 and zoom 6, resulting in 0.374 s per frame. Laser power (Ar 488/35 mW) was set to 80% and 4% was used for imaging. After two pre-bleach images, a circle with a diameter of 4 µm in the nucleus was bleached with 100% laser power with the zoom in option. The recovery was followed as fast as possible for the first 30 s and then at 2 s intervals. The data was analyzed by setting the pre-bleach values to 1 and by producing a linear fit of the first data points to get the import rate.

### RNA-seq

For RNA-seq, PC6.3 cells were plated to 6 well cell culture plates and differentiated for 0-5 days. Total RNA was extracted with a Nucleospin RNA kit from Macherey-Nagel according to the manufacturer’s protocol from quadruplicates. Libraries were prepared for Illumina NextSeq 500 using Ribo-Zero rRNA Removal Kit (Illumina) and the NEBNext Ultra Directional RNA Library Prep at the Biomedicum Functional Genomics Unit (FuGU) according to the manufacturer’s protocols. RNA-seq data sets were aligned using TopHat2 (using Chipster software) to version Rnor_6.0 of the Rattus norvegicus genome with the default settings. Counting aligned reads per genes were performed with HTSeq. Differential expression analysis and Principal components analysis (PCA) were performed with DESeq. List of the transcribed genes was based on cutoff >10 of baseMean parameter for aligned reads counts of all RNA-seq. Gene ontology was performed using DAVID 6.8. RNA-seq data will be available at Gene Expression Omnibus with accession number GSExxx.

### RT-qPCR

PC6.3 cells were plated on 6 well cell culture plates at a density of 80 000 cells per well and differentiated with NGF for 0-8 hours or for 0-5 days. RNA was extracted using the NucleoSpin RNA kit (Macherey-Nagel) according to the manufacturer’s instructions. cDNA was made from 250 ng of RNA using Maxima First Strand cDNA Synthesis Kit for RT-qPCR (Thermo Scientific) and 1:10 diluted cDNA was used for qPCR. qPCR was done with SensiFAST SYBR No-ROX Kit master mix (Meridian Bioscience) using the following primers:

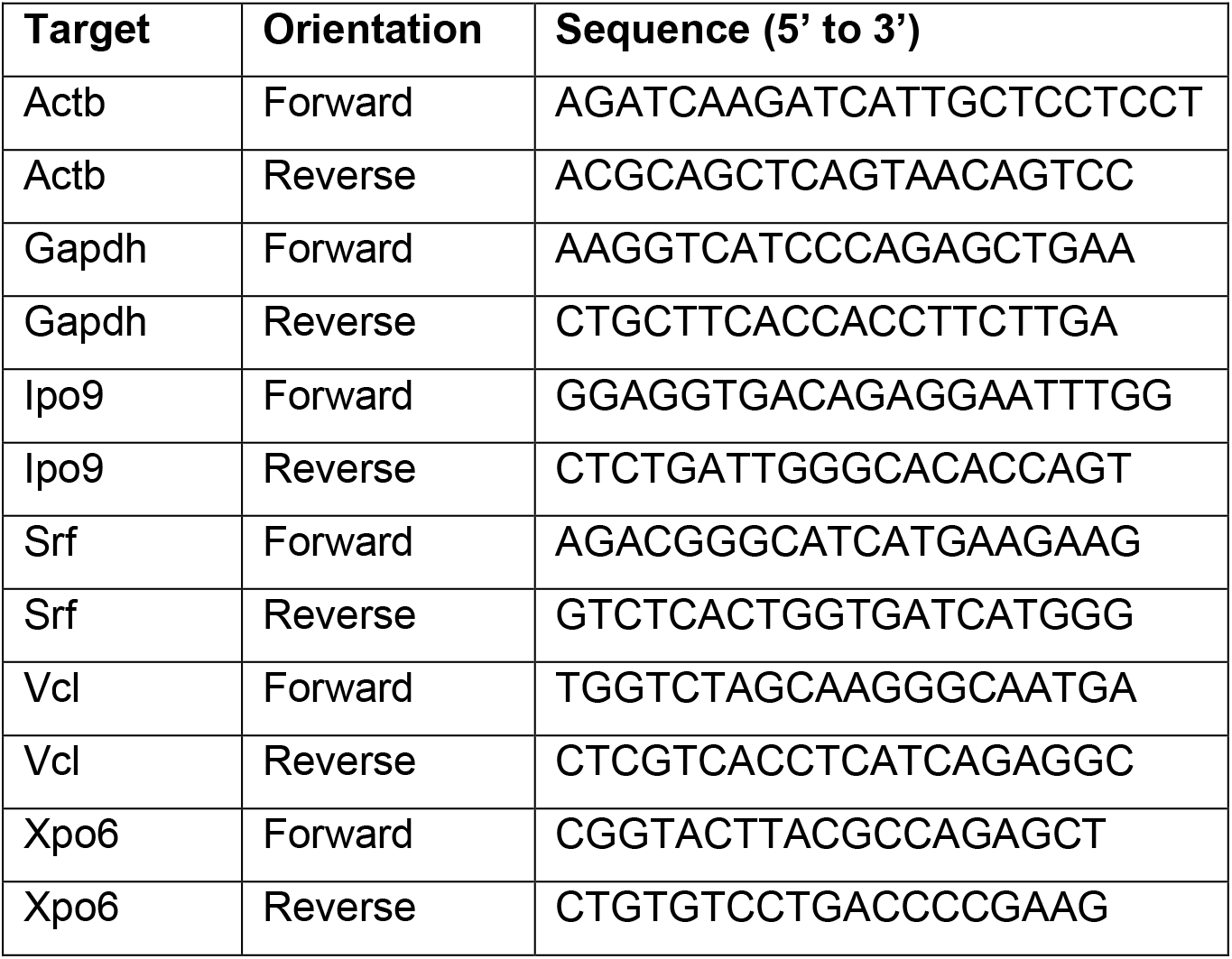

Relative expression levels were calculated by the comparative Ct method, normalizing to Gapdh.

### Statistical analyses

Statistical analyses were performed in Origin 2021b. Normality was tested and when data conformed to a normal distribution, two-tailed one-sample or two-sample t-test was performed. When data was not normally distributed, a two-tailed Mann-Whitney test was used. All statistical tests were done with the significance level of 0.05. The used statistical test is indicated in the figure legends.

## Supplementary figure legends

**Figure S1.**
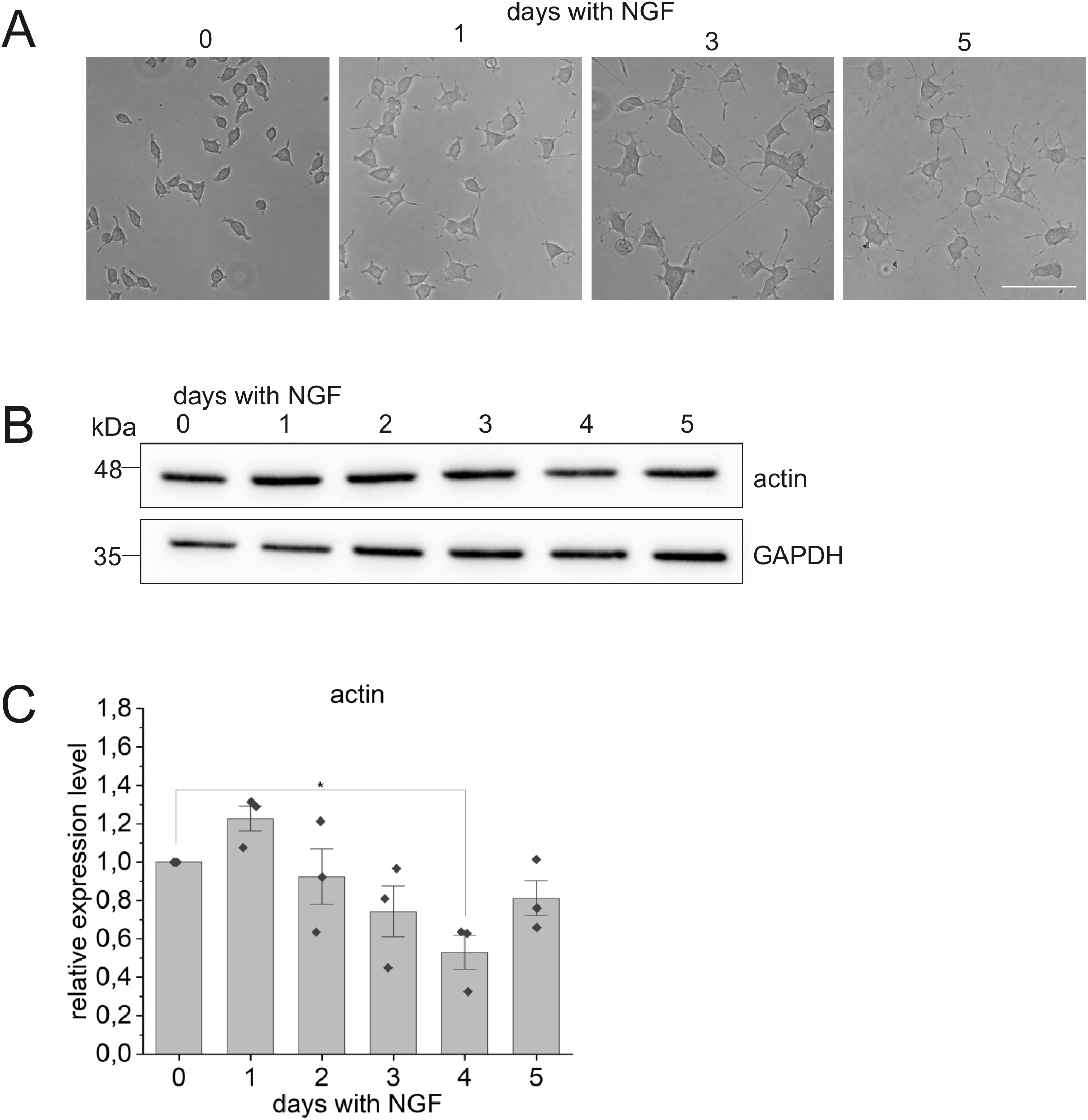
related to figure 1. Images of differentiating PC6.3 cells and assessment of total actin levels. A. Relief phase images of PC6.3 cells during days 0, 1, 3 and 5 of differentiation taken using FLoid Cell Imaging Station. Scale bar 100 μm. B Western blotting of total actin levels during PC6.3 cell differentiation at indicated days after addition of NGF using anti-actin AC-40 antibody that recognizes all actin isoforms and GAPDH as a loading control. Molecular weights on the left. C. Quantification of total actin levels from Western blots. Data is mean (N=3), normalized to undifferentiated sample. Error bars SD, individual data points shown as black dots. Statistically significant differences (P<0.05) with a one-sample t-test: 0 vs. 1 day P=0.096, 0 vs. 2 days P=0.692, 0 vs. 3 days P=0.234, 0 vs. 4 days P=0.045, 0 vs. 5 days P=0.216.

**Figure S2.**
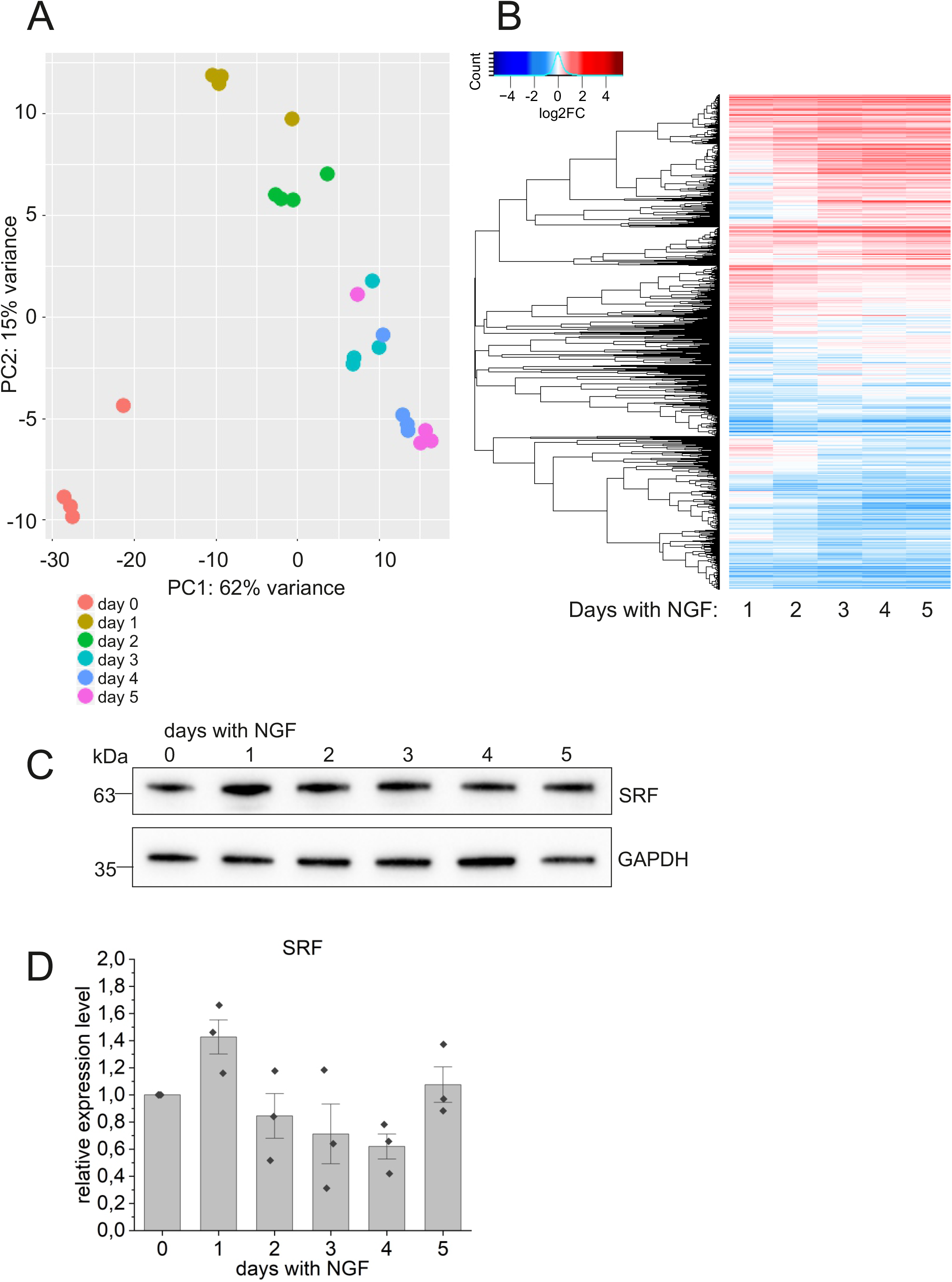
related to figure 2. RNA-seq shows changes in gene expression from the first day of differentiation. A. Principal component analysis results of all RNA-seq samples. Each point represents an RNA-seq sample. Sample groups are indicated by using different colors as indicated in the legend. B. Heatmap showing relative expression of genes (N=12872; see also supplementary table S1) during five days of differentiation following NGF treatment. C. Western blotting of total SRF protein levels during PC6.3 cell differentiation at indicated days after addition of NGF using SRF antibody and GAPDH as a loading control. GAPDH is the same as shown in Fig 4A. Molecular weights on the left. D. Quantification of total SRF levels from Western blots. Data is mean (N=3), normalized to undifferentiated sample, Error bars SD, individual data points shown as black dots. Statistical significance with a one-sample t-test: 0 vs. 1 day P=0.099, 0 vs. 2 days P=0.501, 0 vs. 3 days P=0.375, 0 vs. 4 days P=0.070, 0 vs. 5 days P=0.668.

**Figure S5.**
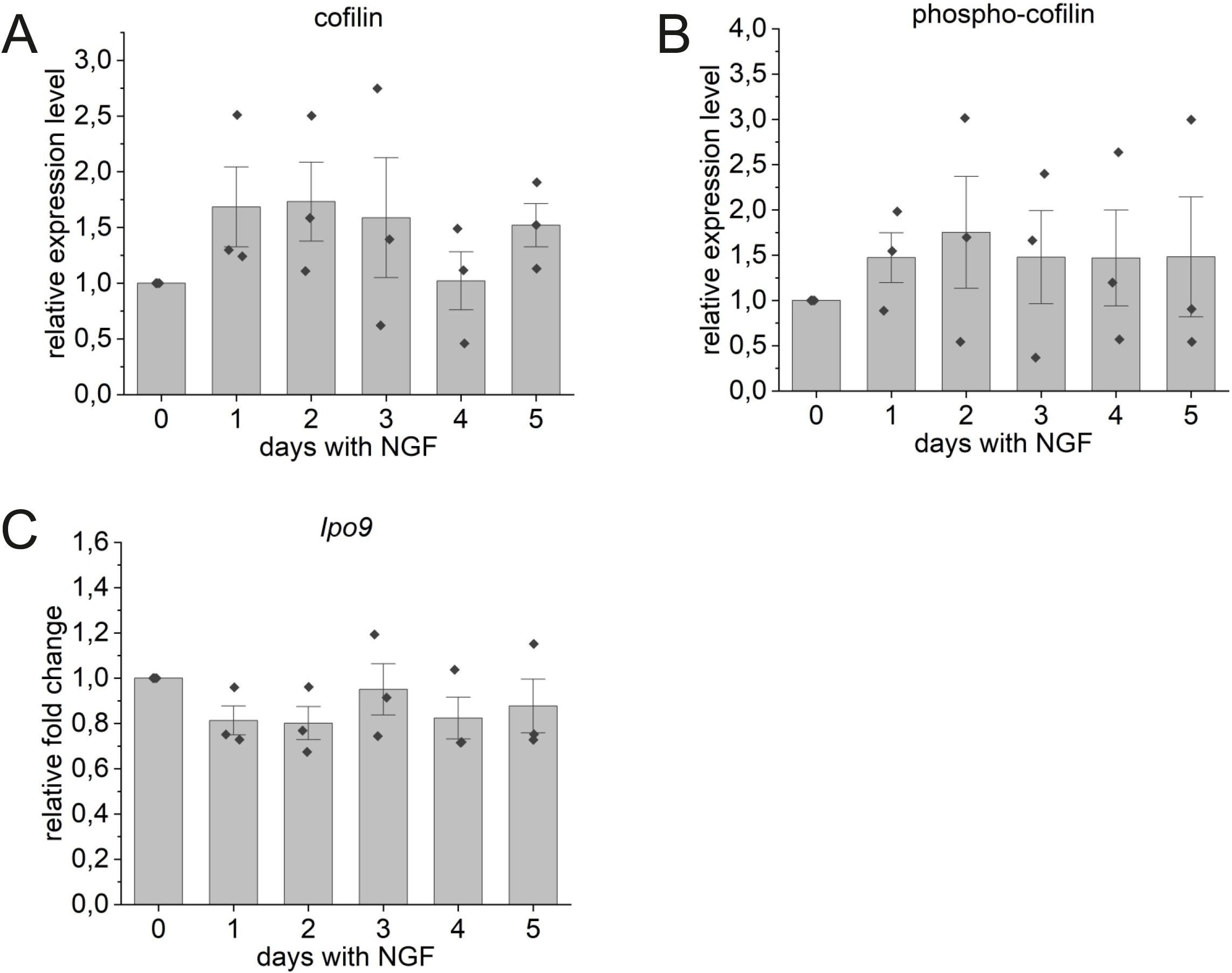
related to figure 5. Protein and mRNA levels of factors required for actin nuclear import remain constant during PC6.3 cell differentiation. A. Quantification of total cofilin levels from Western blots during PC6.3 cell differentiation at indicated days after addition of NGF (example blot shown in Fig 5C). Data is shown as mean (N=3), normalized to undifferentiated sample, error bars SD, individual data points shown as black dots. Statistical significance with a one-sample t-test: 0 vs. 1 day P=0.240, 0 vs. 2 days P=0.215, 0 vs. 3 days P=0.444, 0 vs. 4 days P=0.951, 0 vs. 5 days P=0.145. B. Quantification of phospho-cofilin Western blots, data shown as in S5A. Statistical significance with a one-sample t-test: 0 vs. 1 day P=0.277, 0 vs. 2 days P=0.403, 0 vs. 3 days P=0.505, 0 vs. 4 days P=0.524, 0 vs. 5 days P=0.593. C. qPCR of Ipo9 during PC6.3 cell differentiation at indicated days after addition of NGF. Data is shown as in A. Statistical significance with a one-sample t-test: 0 vs. 1 day P=0.127, 0 vs. 2 days P=0.143, 0 vs. 3 days P=0.742, 0 vs. 4 days P=0.241, 0 vs. 5 days P=0.465.

## Supplementary tables

**Supplementary table S1.** RNA-seq data for all 12872 expressed genes (baseMean expression levels above 10) from day 1 till day 5. DESeq2 analysis of differential expression compared to undifferentiated cells, log2FC values are marked bold.

**Supplementary table S2.** List of up- and down-regulated genes during PC6.3 cell differentiation comparing undifferentiated to 1 day differentiated cells (down1day and up1day) and 1 day differentiated to 5 day differentiated cells (down5vs1 and up5vs1) with gene ontology analysis and MRTF-A target genes indicated.

**Supplementary table S3.** List of MRTF/SRF target genes that display differential expression (Log2FC> 0.6 or <-0.6) at any day of differentiation compared to undifferentiated cells with gene ontology analysis.

---

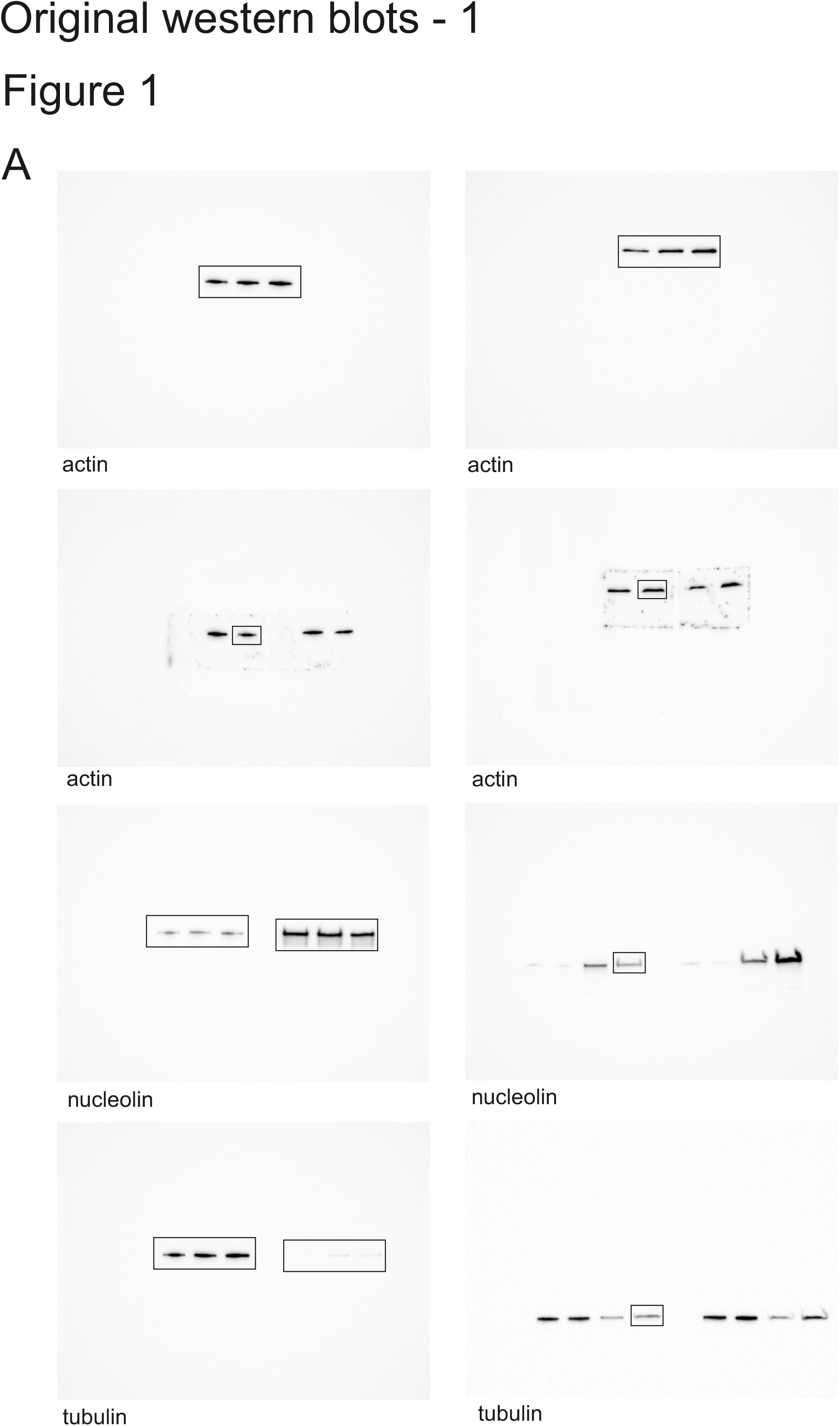

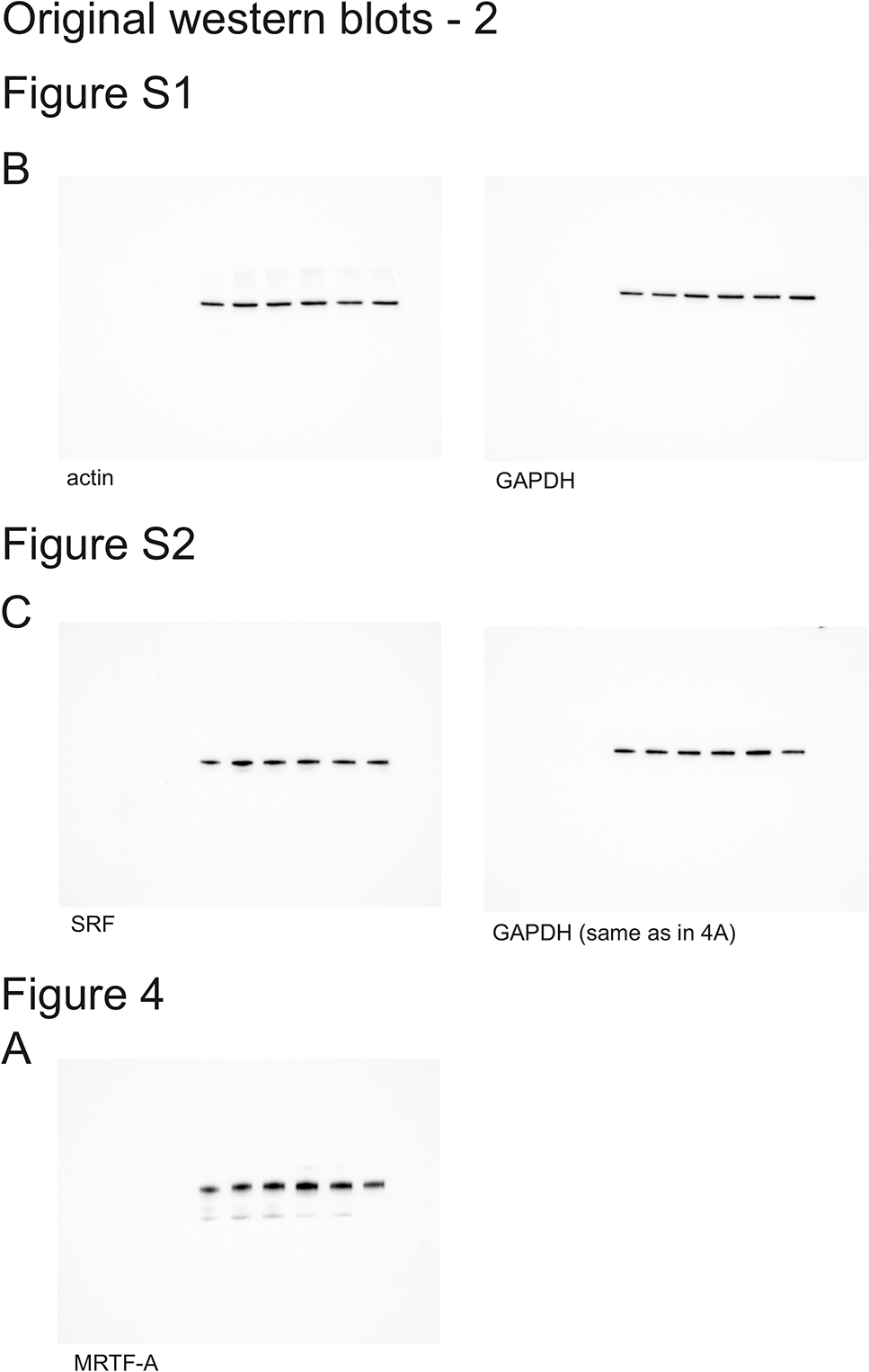

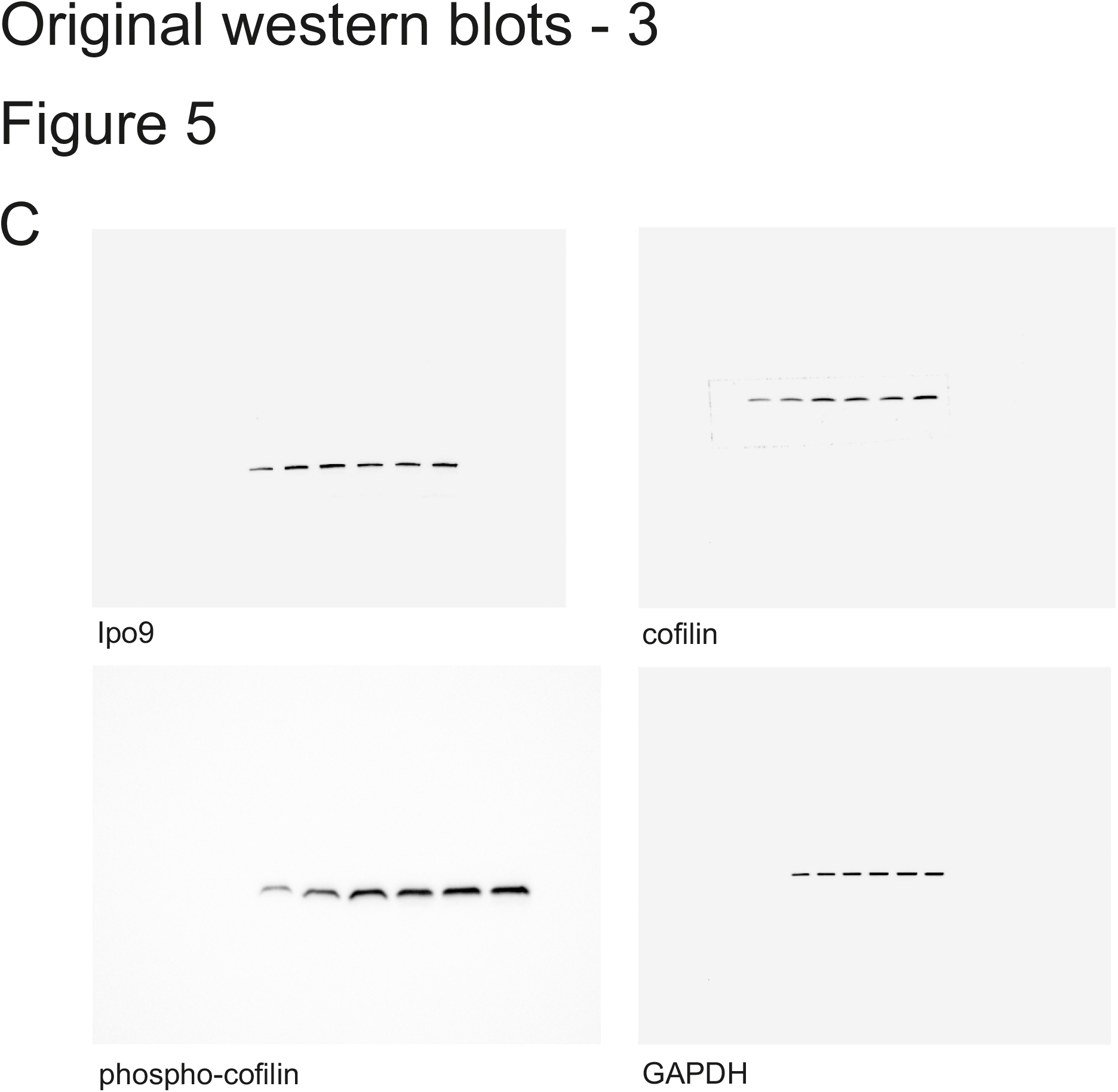

